# A mark-release-recapture approach to characterize tick questing preferences and dispersal distance

**DOI:** 10.64898/2026.05.26.728002

**Authors:** Sabrina Gobran, Jake Brisnehan, Jon Wegryn, Elizabeth Hemming-Schroeder

**Affiliations:** Center for Vector-borne Infectious Diseases (CVID), Department of Microbiology, Immunology and Pathology, Colorado State University, Fort Collins, CO 80523-1685, USA

**Keywords:** tagging, tracking, tick distribution, tick dispersal, field sampling, monitoring, wildlife surveys, mark-recapture

## Abstract

1. Studying tick behavior is crucial for understanding how climate, disturbances, and land-use changes shape tick populations and tick-borne disease risk. Mark-release-recapture studies can provide valuable answers to questions regarding tick movement and behavior, population sizes, and survivorship.
2. Standard tick mark-release-recapture provides limited resolution to understanding individual behaviors, limiting our ability to answer questions that require repeated observations of the same individuals. We developed a new, operationally simple method to track large populations of individual ticks over space and time.
3. We found non-random movement patterns, including directed movement towards grass, vegetation-dependent dispersal distances and rate, and sex-based differences in movement.
4. ***Practical implication:*** This method can be applied to other tick species to assess tick longevity, determine dispersal ranges and rate, and analyze questing behavior and success.

## 1 INTRODUCTION

Tick-borne diseases (TBDs) constitute a greater threat to human health within the contiguous United States than mosquito or flea-borne viruses, bacteria, and parasites (Rosenberg et al., 2018). From 2001-2023, vector-borne disease cases more than doubled in the U.S. (CDC, 2024), with TBDs garnering the highest case reports (Rosenberg et al., 2018). The most notable TBD vector in the U.S. is *Ixodes scapularis* (black-legged tick), which is distributed throughout the eastern and midwestern states and is capable of transmitting *Borrelia burgdorferi* (Lyme disease), among other disease-causing microorganisms (Molaei et al., 2022). *Amblyomma americanum* (Lone star tick), *Amblyomma maculatum* (Gulf Coast tick) and *Dermacentor variabilis* (American Dog tick) also share distribution throughout the eastern U.S. and are capable of transmitting pathogens including *Ehrlichia chaffeensis* (ehrlichiosis), *Rickettsia parkeri* (rickettsiosis), and *Rickettsia rickettsii* (Rocky Mountain spotted fever) respectively (CDC, 2025). Vectors of *Rickettsia rickettsii* also include *Rhipicephalus sanguineus* (brown dog tick), which is distributed ubiquitously throughout the contiguous U.S., as well as *Dermacentor andersoni* (Rocky Mountain wood tick), which is distributed throughout western North America. Finally, *Ixodes pacificus* (western black-legged tick) is the primary vector along the pacific coast, and is capable of transmitting *Borrelia burgdorferi*, *Borrelia miyamotoi* (hard tick relapsing fever), and *Anaplasmosis phagocytophilum* (anaplasmosis) (Eisen et al., 2024). The geographic distribution and incidence of TBDs in the U.S. has been increasing, alongside growing tick populations. These patterns have been attributed to climate warming, reforestation efforts, and human encroachment into tick habitat (Eisen & Eisen, 2023).

Hard ticks (Acari: Ixodidae) are obligate hematophagous arthropods, requiring a blood meal at each life stage to progress to the next developmental state and reproduce (Mans & Neitz, 2004). The majority of hard ticks are three-host ticks, requiring a distinct bloodmeal host at each life stage (Kolonin, 2007). There are two primary strategies that ticks employ for host seeking: ambush and hunting. Hunter ticks actively pursue their hosts by running across the ground, while ambush ticks find their hosts by questing, a behavior where the tick ascends vegetation with their legs extended, waiting for a host to pass (Fourie et al., 1993; Leal et al., 2020; Ramos et al., 2017). To avoid desiccation, ticks will retreat to the soil layer to replenish moisture. Ticks are therefore reliant on their environment for survival, host seeking, and reproduction, and will be impacted by climate and landscape change. It is thus necessary to study tick behavioral patterns within a changing environment to understand and predict TBD risk.

Mark-release-recapture (MRR) is a commonly used tool in ecological research to study animal movement, estimate population sizes, and quantify survivorship. This technique has been used in numerous pest insect and vector-borne disease systems to aid in surveillance and management programs. For example, fluorescent dust is commonly used to study mosquito population sizes, dispersal behavior, insecticide resistance patterns, and disease transmission (Cianci et al., 2013; Russell et al., 2005). Radio telemetry has been used in kissing bugs (*Triatoma* spp.) to assess non-flight dispersal distances and uncover cryptic resting habitats (Hamer et al., 2018). Protein self-marking has been validated in *Culex* mosquitoes and bed bugs (Stuart et al., 2022; Sivakoff et al., 2016). In ticks, MRR has been used to estimate population sizes and survival probabilities, determine dispersal ranges and direction, and identify preferred habitats (Buczek et al., 2017; Carroll et al., 1991; Carroll & Schmidtmann, 1996; Goddard, 1993; James et al., 2022; Simmons et al., 2015).

Although tick MRR has provided insight into tick behavior, there are limitations to the methods commonly employed. Most MRR studies in ticks involve batch marking with either fluorescent powders or paint applied to the scutum or legs to identify ticks collected and released within a certain timeframe. Resolution to the individual level is impossible following a batch marking strategy. Additionally, fluorescent powders or dust can be short-lived, providing uncertainty to whether ticks have been previously marked and leading to distorted population estimates. Here, we designed an effective system to uniquely mark ticks using nail polish so that large numbers of ticks can be marked and individually identified upon innumerable capture events. This method is longer lasting than powder or dust marking and is effective for marking large populations using a three-color system. Additionally, this method does not require the extreme dexterity required to paint unique patterns or numbers onto the scutums of small-bodied ticks. Altogether, this approach allowed us to precisely track individual tick movement over space and time, which can be used to assess tick longevity, determine dispersal ranges and rate, and analyze questing behavior patterns.

In this study, we validated our MRR approach by quantifying *D. andersoni* dispersal distances and vegetation preferences in Northern Colorado. We constructed five continuous 10 m^2^ study plots in Rustic, Colorado, which were surveyed by active dragging 1-3 times per week for eight weeks. Ticks were marked with nail polish using a unique three-color system and released at predetermined vegetation sites representing the overall diversity of plants in the area. Recaptured ticks were identified based on their unique markings. We recorded the vegetation type each tick was collected from and the straight-line distance to its prior release site. The method described here can be applied to other tick species, providing the potential to increase our understanding of arthropod behavior at an individual level.

## 2 MATERIALS AND METHODS

### 2.1 Study Site

This study was conducted across five study plots at one site with known moderate-to-high *D. andersoni* presence in the Poudre Canyon located in Larimer County, Colorado (Figure 1). The site is relatively flat and situated at approximately 2,200 m of elevation, bordering the Cache la Poudre river on one side. The climate is characterized by cold winters and hot, dry summers. The vegetation consists of mixed grasses, shrubs, ponderosa pine (*Pinus ponderosa*), Douglas fir (*Pseudotsuga menziesii*), and common juniper (*Juniperus communis*). Predominant grass and shrub types include blue grama (*Bouteloua gracilis*), cheatgrass (*Bromus tectorum*), smooth brome (*Bromus inermis*), big sagebrush (*Artemisia tridentata*), and antelope bitterbrush (*Purshia tridentata*). Although this study specifically targeted *D. andersoni* ticks, other tick species including *Dermacentor variabilis* and *Dermacentor albipictus* may be found in this region (Hutcheson et al., 2021).

**Fig. 1.**
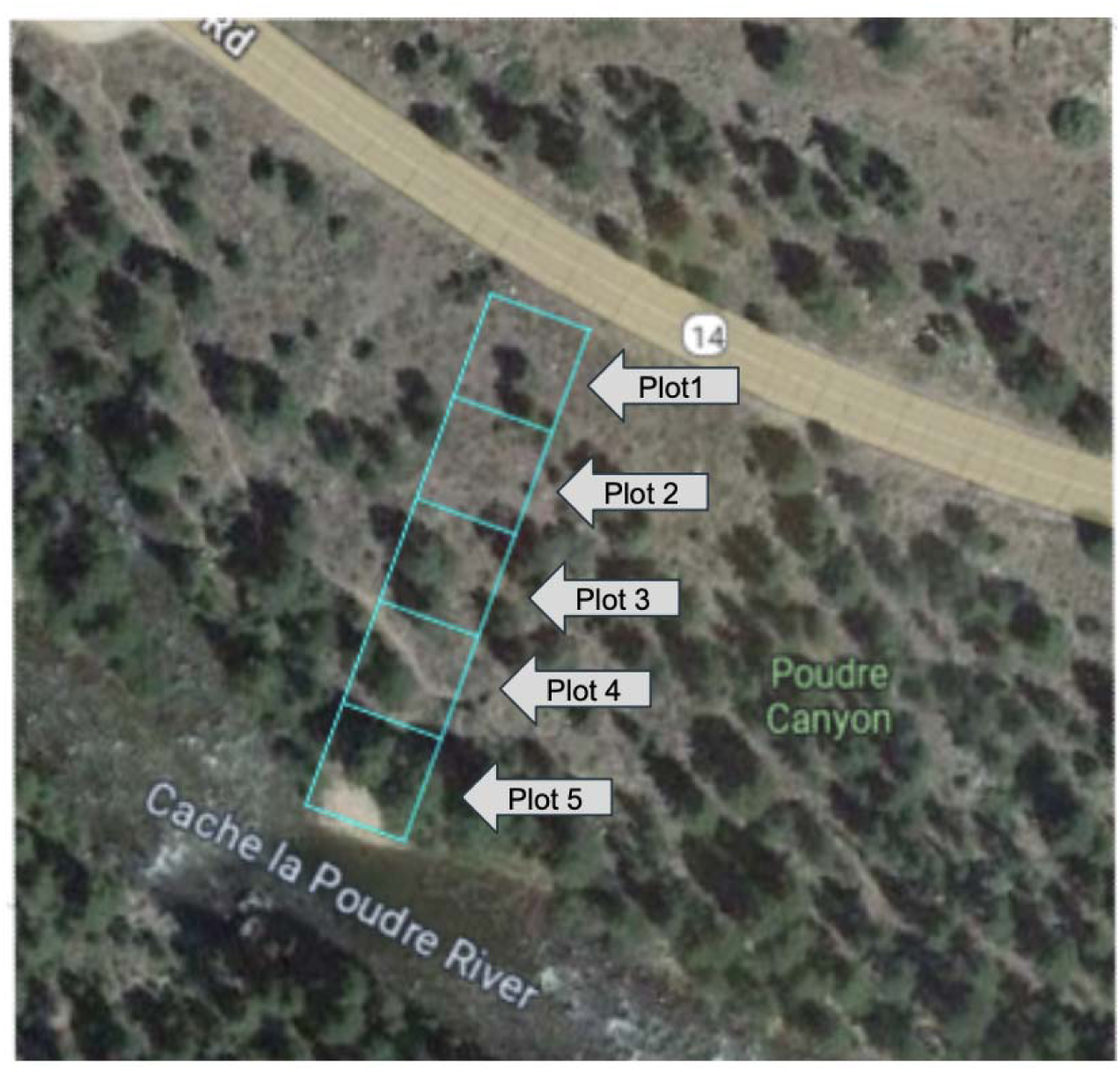
Plot design of tick sampling site.

Vegetation data was collected from each plot using the Line-Intercept sampling technique to identify vegetation types and estimate their abundance within the five sample plots (Table 1). This technique was utilized by running five parallel transect lines equidistant within each plot, using meter tapes. We recorded the distance along the meter tape that was intercepted by each vegetation type, then identified and sorted it into appropriate categories.

**Table 1.** Vegetation abundance (%) by plot.

| Plant Species | Plot 1 | Plot 2 | Plot 3 | Plot 4 | Plot 5 |
| --- | --- | --- | --- | --- | --- |
| Juniper | 5.6 | 7.4 | 0.9 | 0.0 | 0.0 |
| Litter | 52.5 | 44.9 | 60.0 | 47.5 | 52.9 |
| Bare ground | 8.1 | 6.1 | 2.8 | 5.4 | 5.4 |
| Shrub | 12.0 | 20.0 | 8.8 | 21.0 | 17.2 |
| Grass | 21.7 | 17.1 | 27.5 | 26.2 | 24.1 |
| Sapling | 0.0 | 0.0 | 0.0 | 0.0 | 0.4 |
| Other | 0.0 | 4.6 | 0.0 | 0.0 | 0.0 |

### 2.2 Study Organism

*D. andersoni* are the primary tick species in Northern Colorado responsible for the transmission of Rocky Mountain spotted fever (*Rickettsia rickettsii)*, tularemia (*Francisella tularensis*), and Colorado tick fever virus (Eisen et al., 2007). *D. andersoni* is known to inhabit the Mountain West, with populations ranging from the southern parts of Saskatchewan, Alberta, and British Columbia in the north and from Colorado to eastern California in the south (Bishopp & Trembley, 1945; Burgdorfer, 1969; CDC, 2025). In Colorado, *D. andersoni* abundance peaks between 2,200–2,400 m of elevation, although peak population abundance may occur at lower elevations in colder areas to the north, or at higher elevations in warmer areas to the south (Eisen et al., 2007). *D. andersoni* are not host specific, with immature stages typically feeding on small mammals and adults preferring larger mammals (Bishopp & Trembley, 1945; James et al., 2006; Sonenshine et al., 1976). They find their hosts using an ambush strategy, and as such must utilize vegetation for questing. It remains unknown whether *D. andersoni* exhibit questing vegetation preference, their dispersal patterns, and their non-host mediated dispersal range.

### 2.3 Field Collection

Five continuous 10 m^2^ study plots were established for surveillance and monitored by active flagging/dragging 1-3 times per week from April-June 2025. Whether flagging or dragging was used was determined by suitability to the understory type. Dragging was used for tick collection from grass, leaf litter, and bare ground. Flagging was used for tick collection from shrub, juniper, and sapling. The cloth was checked every time the predominant understory vegetation changed. Attached nymphal and adult ticks were removed using forceps and placed into vials. Larval ticks were discarded. The entire area of the study plots was sampled systematically, as well as the surrounding environment.

Ticks were visually inspected for morphology consistent with *D. andersoni* in the field upon first collection. Life stage and sex was also assessed and recorded before marking. Ticks were marked with Sally Hansen Insta-Dri nail polish (Sally Hansen, Morris Plains, NJ USA) using a unique three color combination adapted from White *et al* (White et al., 2020). This color code consisted of a base coat, a superior dot, and an inferior dot (Figure 2). Six distinct polish colors were used, allowing for 120 unique color combinations. Ticks were immobilized on a ball-bearing with putty and painted with a size 000 insect pin, taking care to avoid the ticks’ eyes and spiracular plates (White et al., 2020). Quick dry nail polish was selected to minimize the time required to mark each tick. The nail polish was allowed to dry between painting the base color and dots and before release. The time to mark each individual tick was between 7 and 10 minutes, including drying time.

**Fig. 2.**
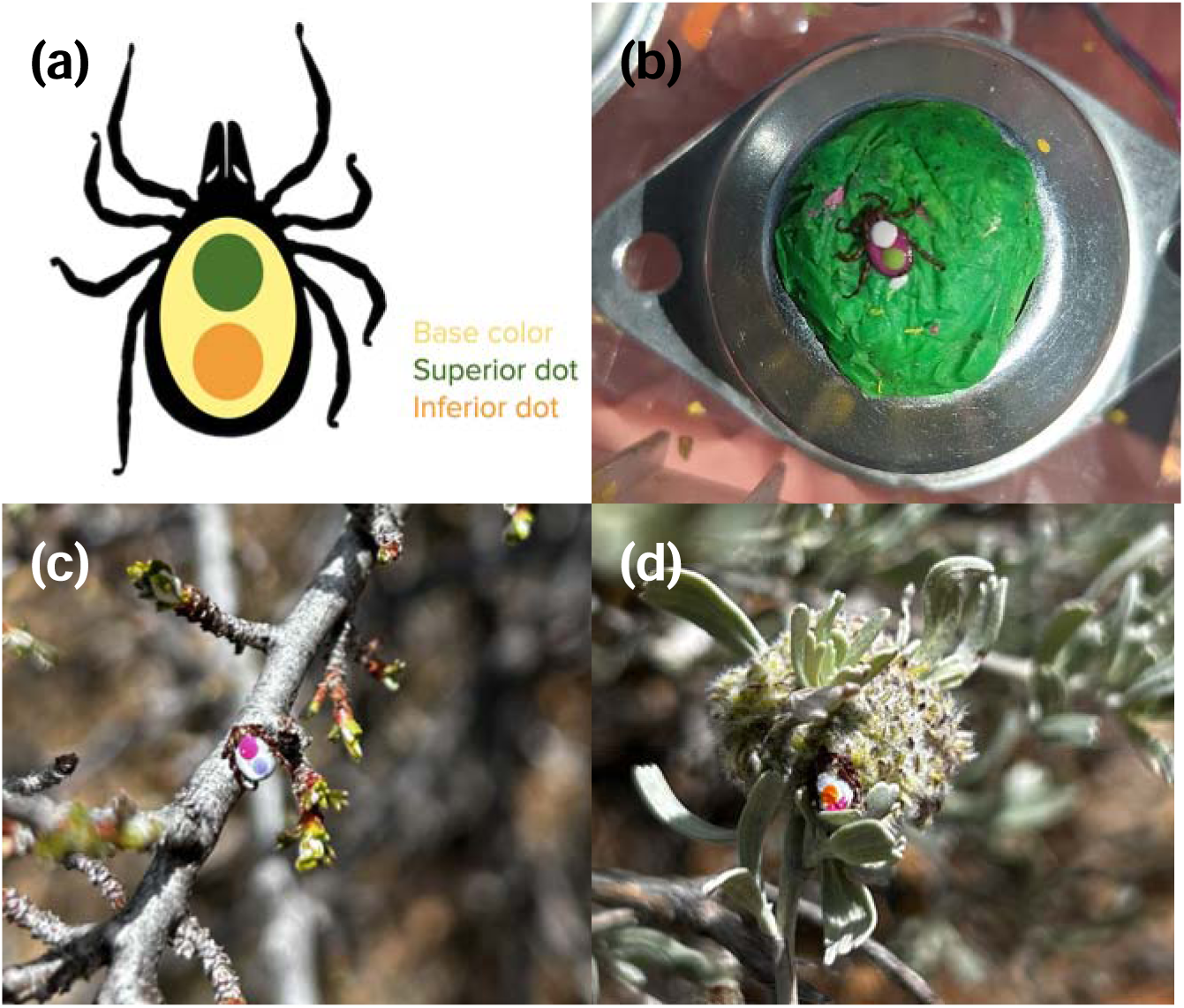
Conceptual marking design (a), example of an immobilized tick after marking (b), and ticks after release onto selected vegetation (c, d).

After marking, ticks were released at designated release points onto selected vegetation types. Release vegetation included bare ground, grass, shrub, juniper, sapling, and leaf litter. All release points were located in plot 3 for its centralized location, except for the sapling release point. Plot five was the only plot containing saplings, and therefore it was selected as the sapling release point.

The vegetation type each tick was collected from was recorded for all ticks collected within the study plots. Upon recapture, the dispersal distance was also recorded, measured as the straight-line distance from the collection location to the release location. Ticks were collected and preserved in DNA/RNA shield (Zymo, Irvine, CA, USA) at -20°C during the final week of sampling for further analyses.

### 2.4 Laboratory

Nail polish residue was removed from the ticks using 10% acetone. Ticks were then sterilized by immersion in 10% bleach, 70% ethanol, and distilled water to remove external parasites and debris. Tick life stage, species, and sex was confirmed in the lab using a dissecting microscope and dichotomous key (Evert E. Lindquist et al., 2016).

### 2.5 Analyses

Analyses were performed in R (version 4.5.1). Dispersal distance and dispersal rate were compared using log-transformed t-tests or Wilcoxon rank-sum tests, depending on whether assumptions of normality were met. Differences in dispersal distance between male and female ticks and among release vegetation types were evaluated using two-sample t-tests and pairwise t-tests, respectively, applied to log-transformed dispersal distance values. Holm-Bonferroni correction was used to adjust for multiple comparisons when conducting pairwise tests across vegetation types. Differences in dispersal rate between sexes and among release vegetation types were assessed using Wilcoxon rank-sum tests, with Holm correction applied for multiple comparisons across vegetation types. To determine whether ticks moved towards certain vegetation types, we visualized how many ticks were released at each vegetation type and how many were collected from each vegetation type. Then we used chi-squared tests with Benjamini, Hochberg adjustment to test for significant associations. We used linear models (i.e., days since last capture, study date) to determine the relationship between dispersal distance and time using the lm() base R function. To visualize plot level movement patterns, the igraph R package was utilized after constructing an adjacency matrix of movements from release sites to capture sites.

## 3 RESULTS

Overall, 50 host-seeking nymphal and adult ticks were collected, marked, and released at the study site. Sampling began in mid-April and continued through the first week of June. Five ticks were collected for the first time during the final week of sampling (06/05/25-06/06/25) and were not re-released. In the field, the 55 ticks were identified as 27 *D. andersoni* females, 25 *D. andersoni* males, 2 *D. andersoni* nymphs, and 1 other *Dermacentor* species. In the final week of collections, 20 ticks were captured and collected for later analysis. Of these, 10 were identified as *D. andersoni* females, 9 were identified as *D. andersoni* males, and 1 was identified as *D. albipictus*. The *D. albipictus* individual was removed from further analyses. All microscopic identifications were consistent with field identifications except one nymph, which was reclassified as an adult female based on the presence of genital aperture. The species and life stage of ticks not collected during the final week of sampling was unable to be confirmed via microscopy.

Final collections occurred 54 days after the initial release. Ticks were present in the study area for the entire duration of the study and ticks were collected on every sampling day. Peak abundance of questing ticks, determined by the largest number of ticks collected on a single sampling day, occurred on both May 17 and May 23, 2025 (Figure 3). By the fifth sampling day the majority of ticks collected were marked and by the tenth sampling day all ticks collected were marked, with the exception of the final sampling day, when there was an even number of marked and unmarked ticks collected.

**Fig. 3.**
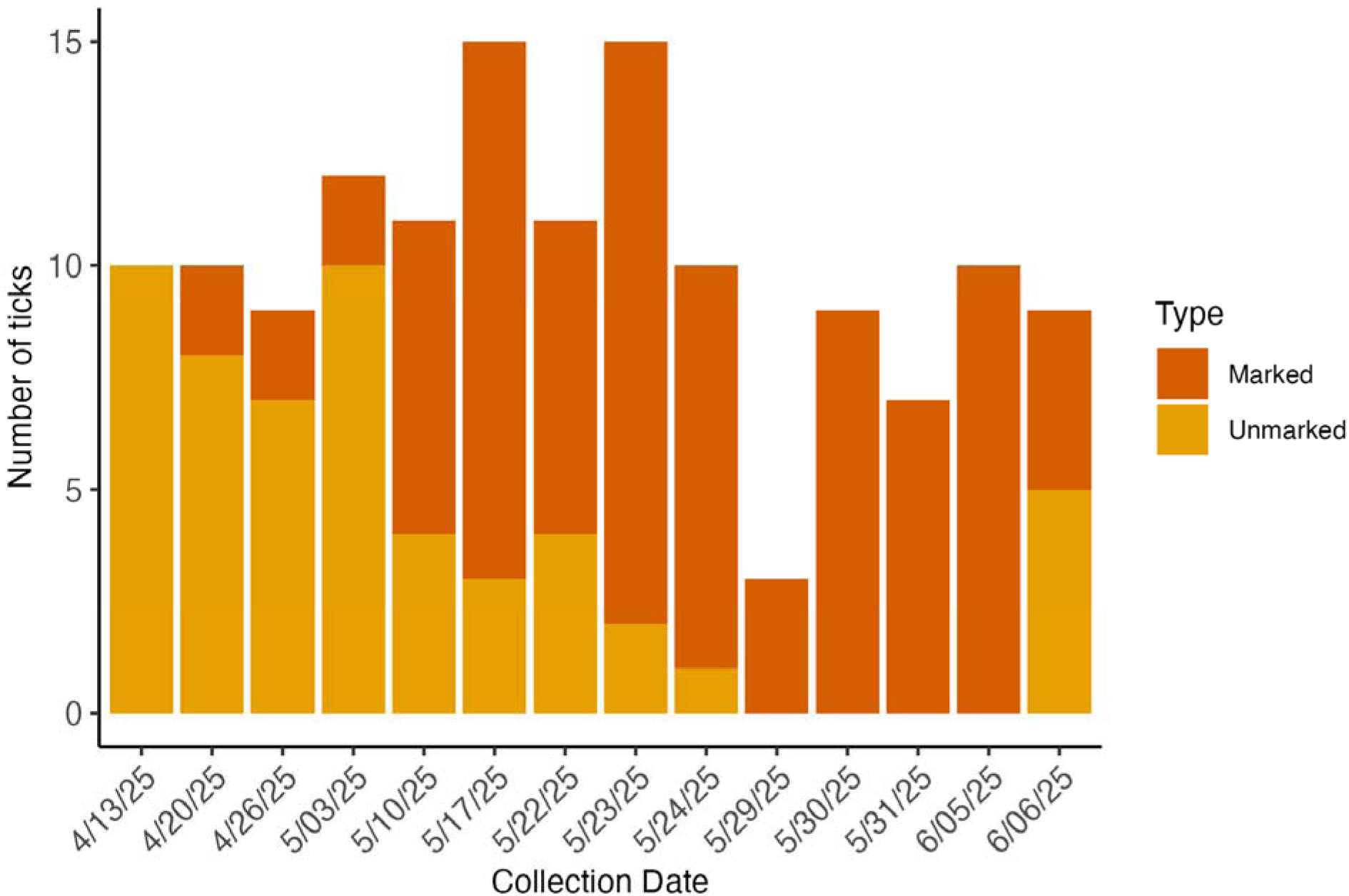
Total numbers of marked and unmarked host-seeking *D. andersoni* ticks by collection date.

On average, ticks were recaptured 2.8 times (SD = 1.5; Figure 4A), with a maximum of six captures of an individual tick (n = 3). Thirteen ticks were released but never recaptured (13/50, 26%). The mean distance travelled between captures was 4.5 m (SD = 4.6; Figure 4B), ranging from no movement (0 m) to 19.1 m. The number of days between captures averaged 10.5 days (SD = 9.1; Figure 4C). More than 80% of recapture events occurred within 2 weeks of the initial release, although the maximum time between captures was 41 days (n = 1). The average rate of movement was 0.67 m/day (SD = 1.1; Figure 4D), with most ticks moving <2 m/day. A small number of individuals (n = 8) exhibited higher movement rates (>2 m/day), with the fastest rate at 6.3 m/day.

**Fig. 4.**
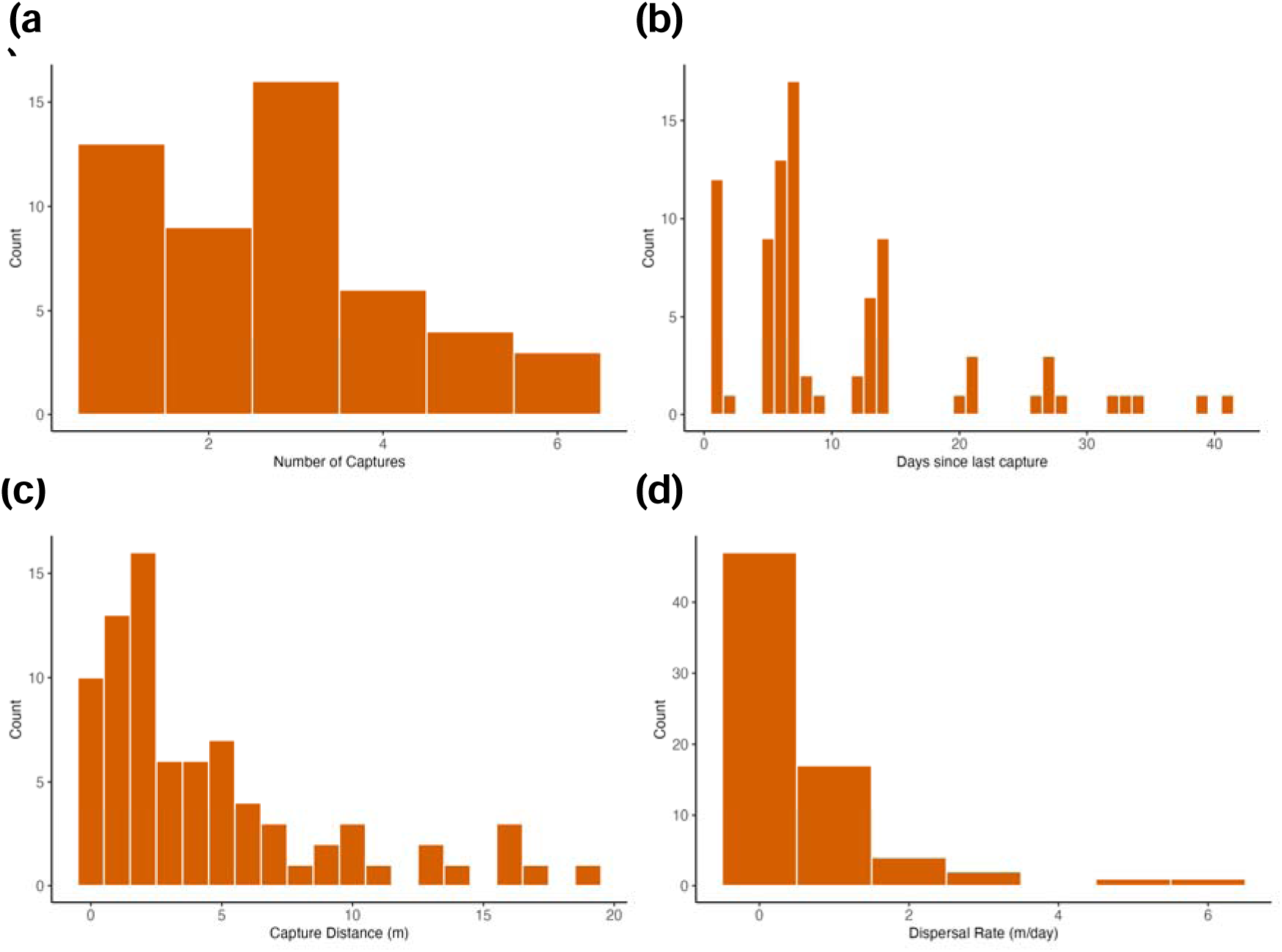
Histograms showing the number of captures per tick (a), the number of days between captures (b), the overall dispersal distance between captures (c), and the dispersal rate (d).

Movement rates varied by sex, averaging 0.85 m/day in adult females and 0.58 m/day in adult males (Figure 5), although this difference was not significant (p = 0.5268, Wilcoxon rank sum test). Using overall distance, adult females moved significantly further than adult males, averaging 6.036 m and 3.567 m, respectively (p = 0.044; t-test).

**Fig. 5.**
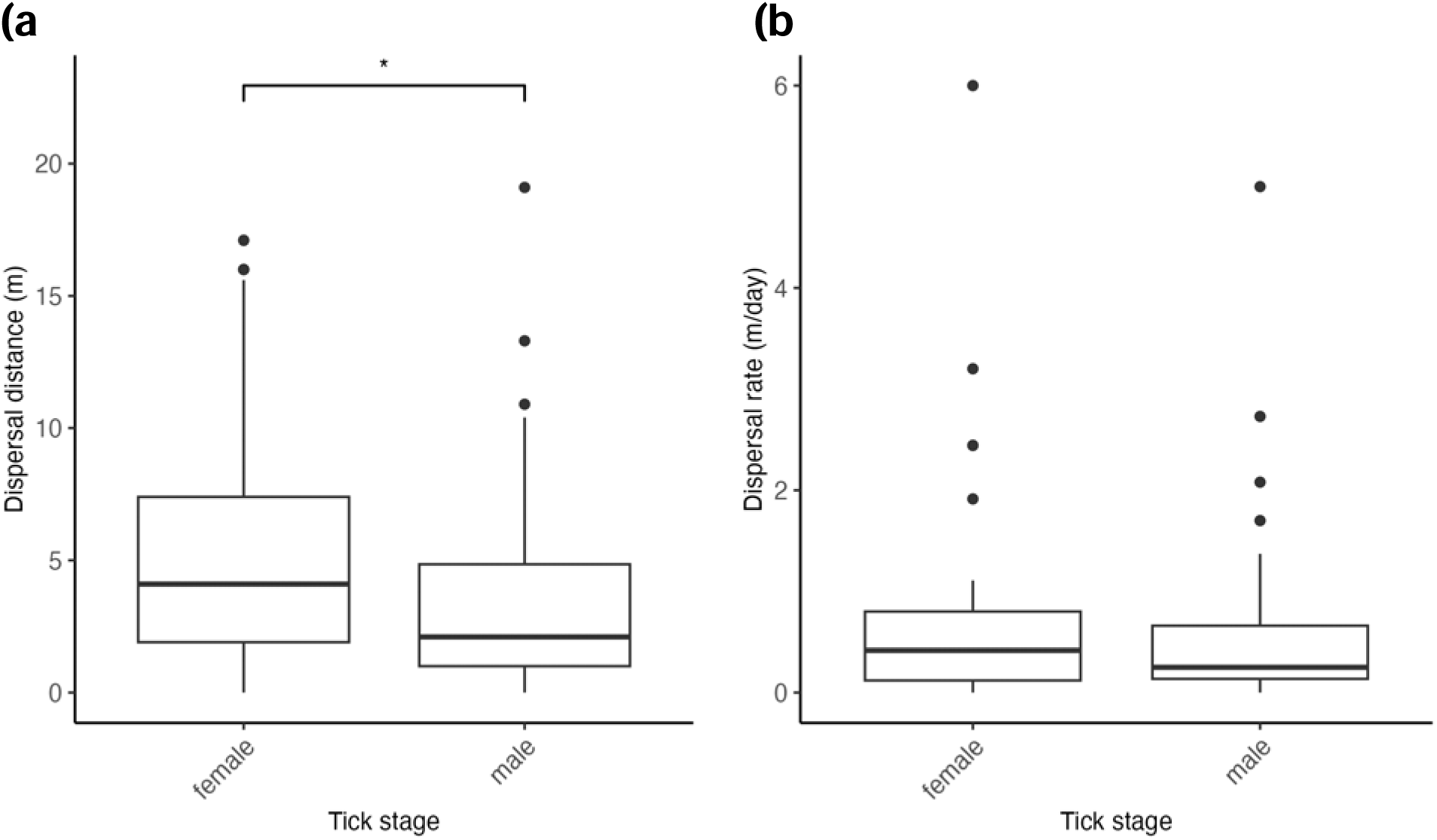
Boxplots of dispersal distance (a) and dispersal rate (b) by life stage.

Ticks dispersed the farthest when placed on leaf litter (7.62 m) or juniper (6.76 m) and ticks dispersed the least amount when placed on grass (1.64 m) or bare ground (2.45 m; Figure 6A). There were significant differences in overall dispersal distance when comparing ticks placed on bare ground and leaf litter (p = 0.016; log-transformed pairwise t-test), bare ground and juniper (p = 0.018; log-transformed pairwise t-test), grass and leaf litter (p = 0.005; log-transformed pairwise t-test), and grass and juniper (p = 0.005; log-transformed pairwise t-test; Figure 6A). Dispersal rates were similarly highest when placed on leaf litter (1.60 m/day) and juniper (1.25 m/day) and lowest when placed on grass (0.22 m/day) or bare ground (0.19 m/day), and there were significant differences in dispersal rate when comparing ticks placed on bare ground and leaf litter (p = 0.025; Wilcoxon rank sum test), bare ground and juniper (p = 0.008; Wilcoxon rank sum test), grass and leaf litter (p = 0.029; Wilcoxon rank sum test), and grass and juniper (p = 0.018; Wilcoxon rank sum test; Figure 6B).

**Fig. 6.**
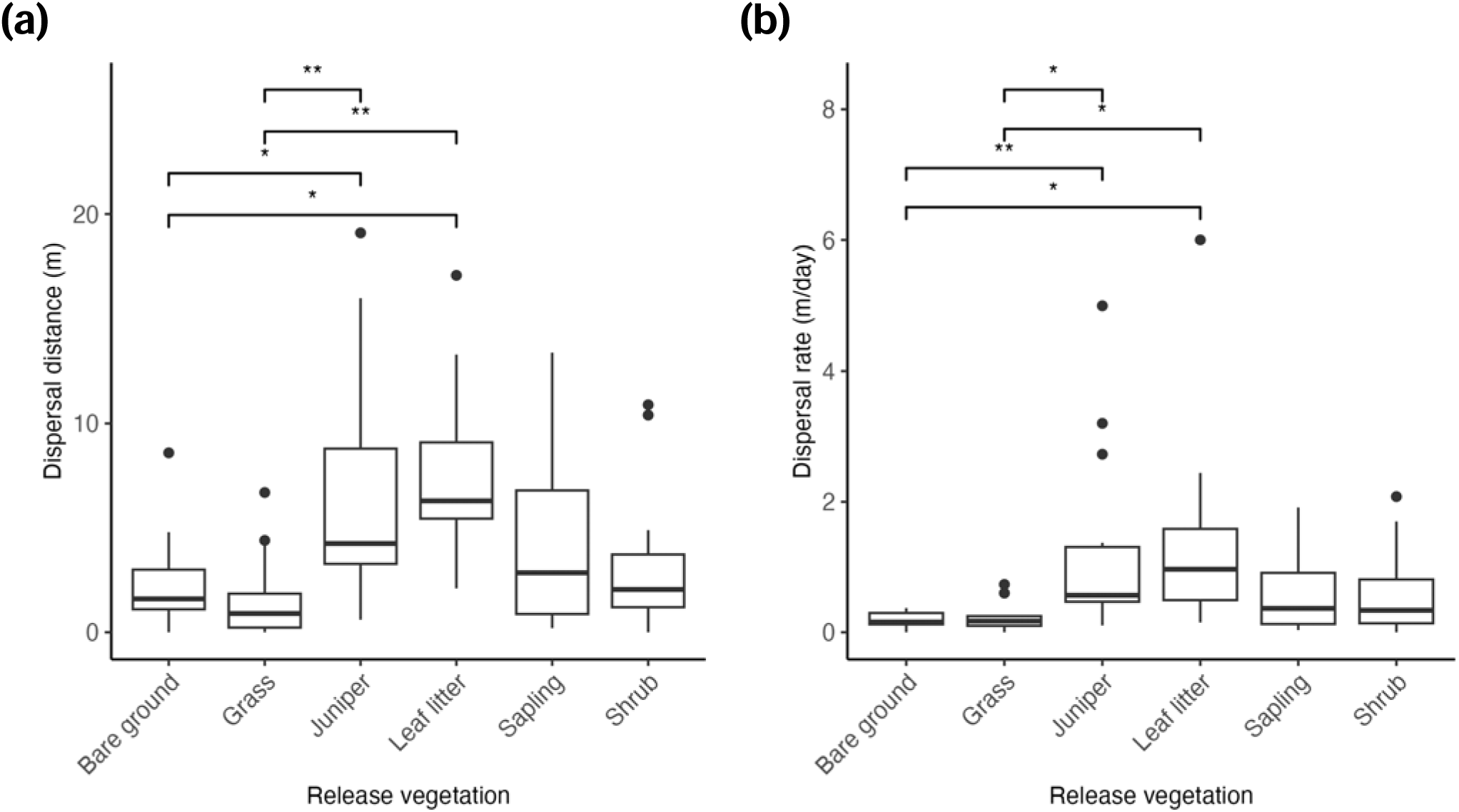
Boxplots of dispersal distance (a) and dispersal rate (b) by release vegetation type.

When comparing the number of ticks captured on vegetation types in relation to the number of ticks released at those vegetation types, we found that ticks were captured at disproportionately higher rates on grass (p = 0.013; Figure 7), whereas ticks were captured at disproportionately lower rates on bare ground (p < 0.001), juniper (p = 0.0013), sapling (p < 0.001), and shrub (p = 0.0013). Ticks were captured on litter at a rate commensurate with the number of ticks released on litter (p = 0.74).

**Fig. 7.**
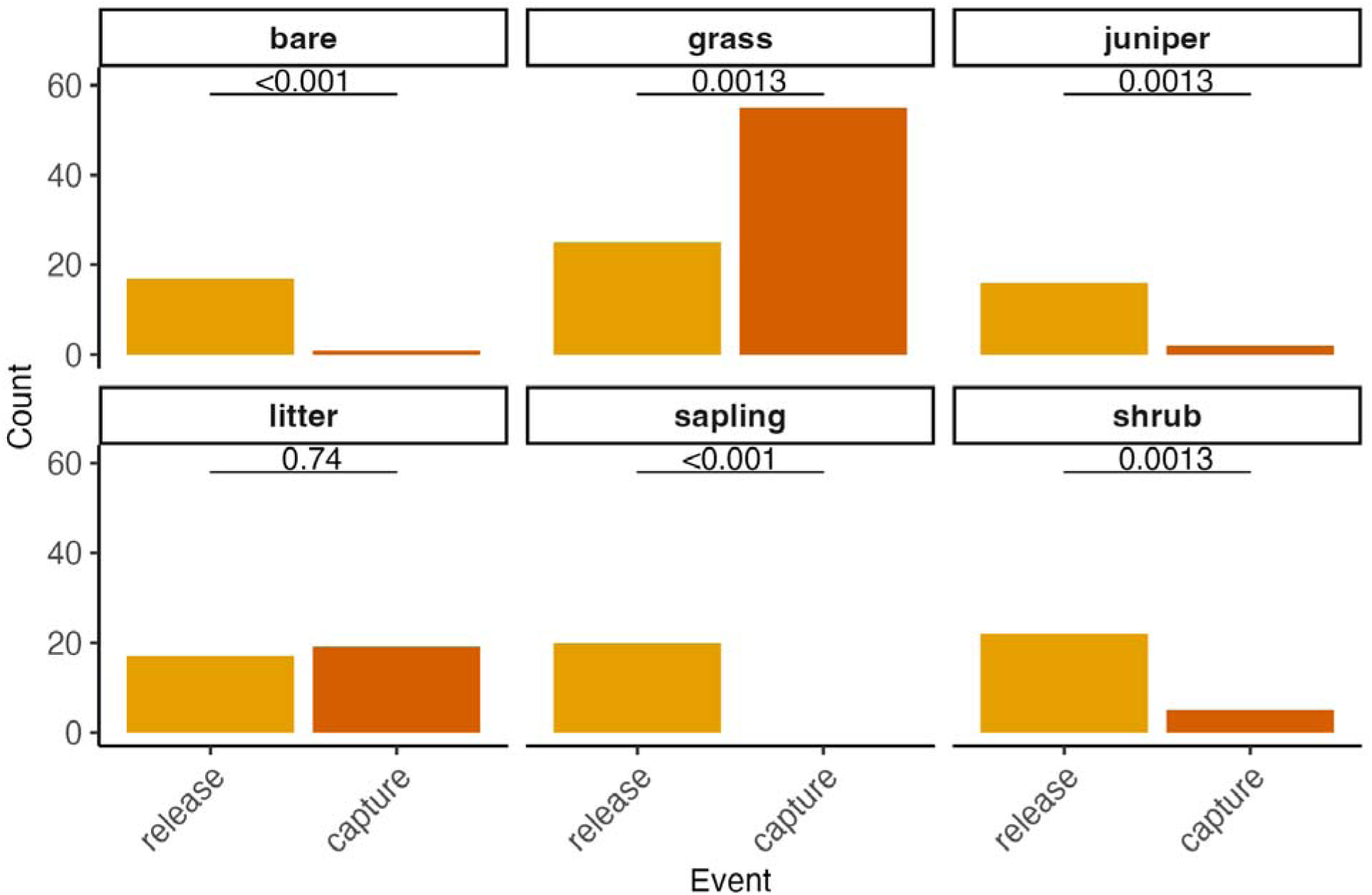
Number of ticks released and recaptured from each vegetation type.

To evaluate if ticks disperse at a constant rate over time, we fit a linear model with distance from last capture as the response and days since last capture as the predictor (Figure 8). Although distance from the last capture increased by 0.122 meters per day (*β* = 0.122), days since last capture explained only 2.6% of the variation in distance (*R²* = 0.026), indicating substantial unexplained variability. The relationship between days since last capture and distance travelled was not significant (p > 0.1).

**Fig. 8.**
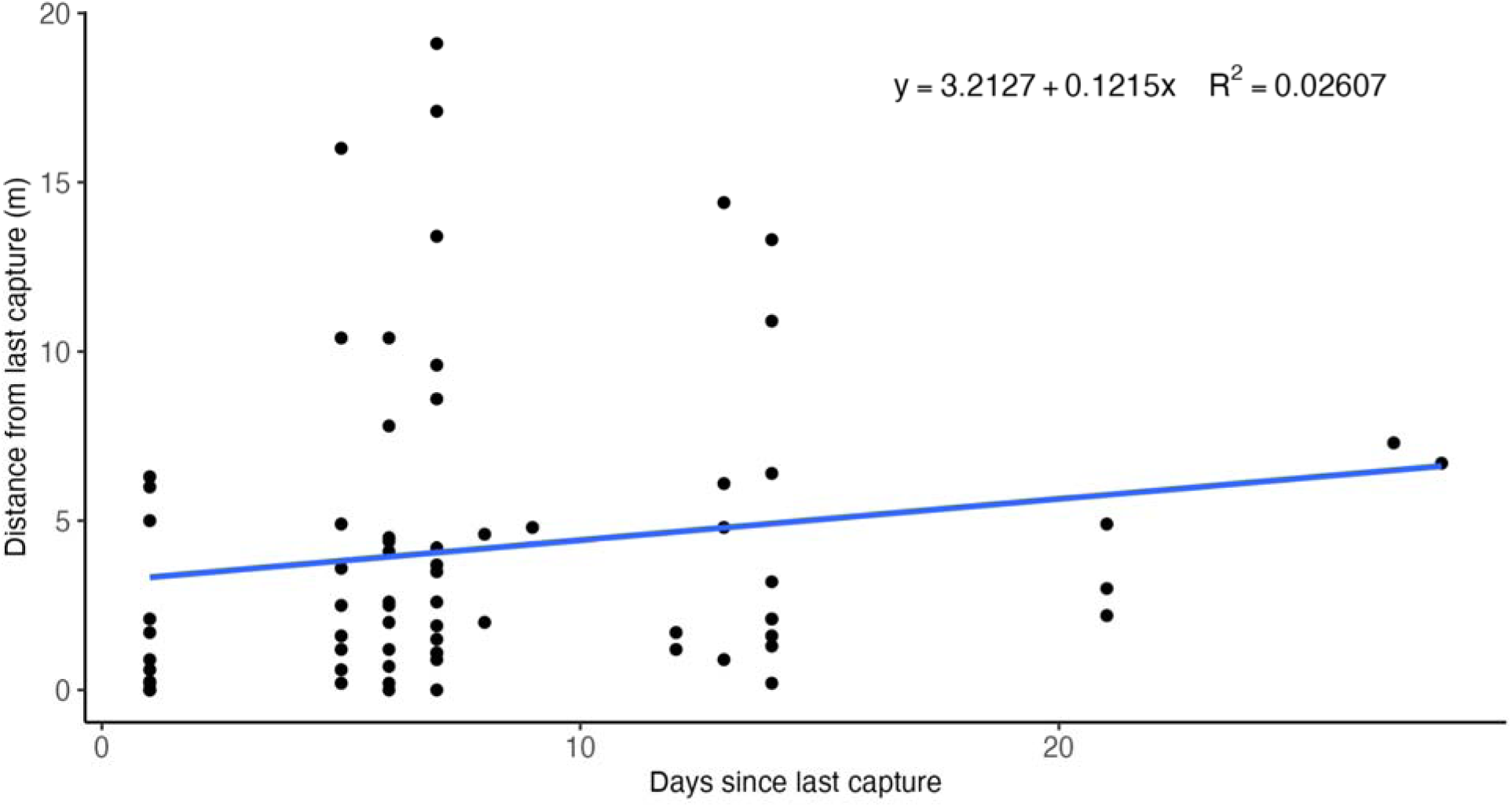
Relationship between dispersal distance and time since last capture.

To evaluate how dispersal changes across the season, we fit a linear model with distance from last capture as the response and study date as the predictor (Figure 9). Each change in calendar date corresponded with a decrease in 0.150 m travelled (*β* = -0.150) and study date explained 6.9% of the variation in distance (*R²* = 0.069). The relationship between study date and distance travelled was significant (p = 0.027).

**Fig. 9.**
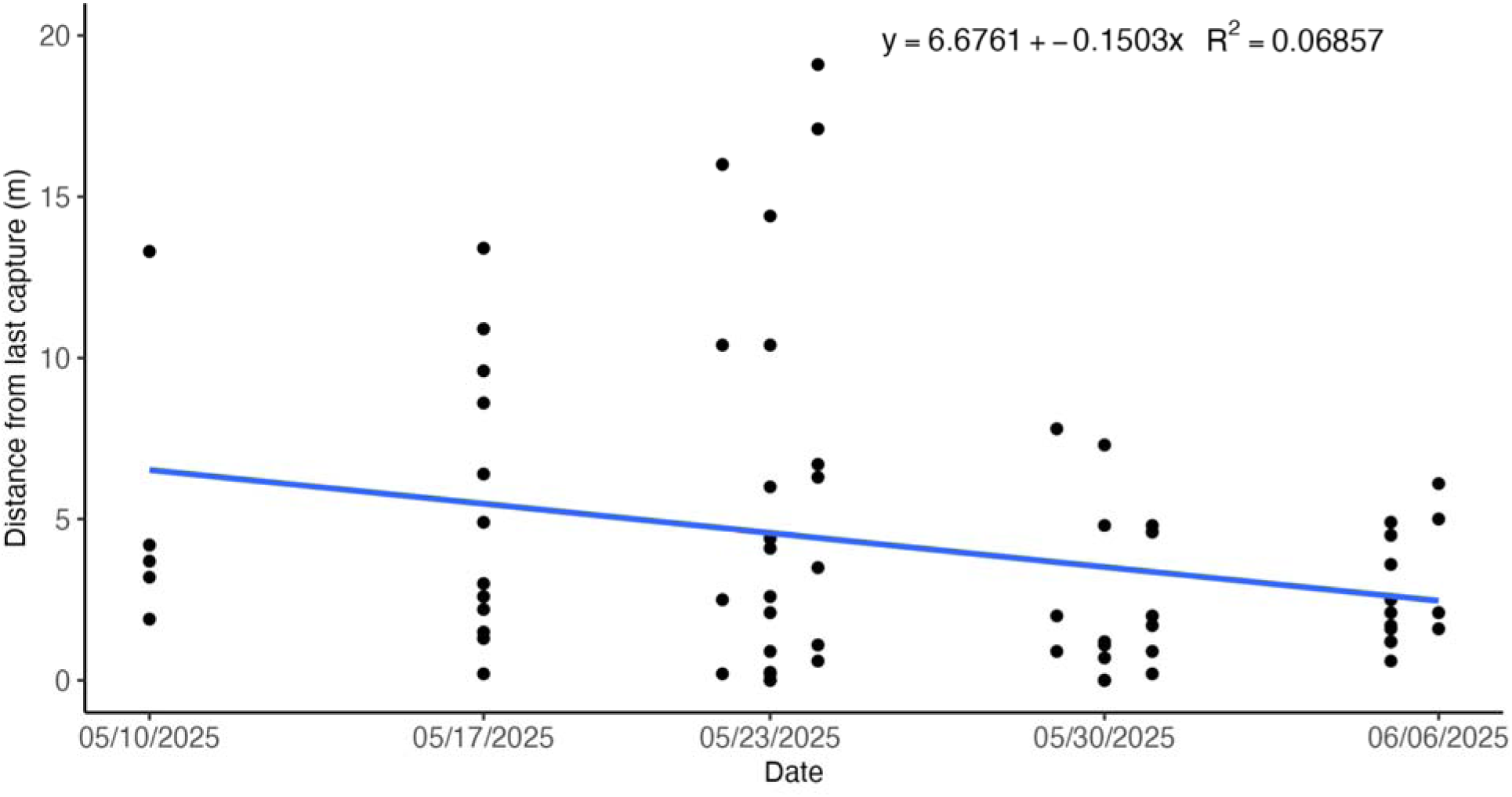
Relationship between study date and dispersal distance.

To best visualize the movement of individual ticks from the release plot to their next capture, we created an adjacency matrix to use for a directed graph (Figure 10). Each arrow shows the movement of ticks within and across plots. Arrows that start and end in the same plot represent ticks that were released and captured within the same plot boundaries. Because ticks were only released in plots 3 and 5, arrows are only shown originating in those plots. Arrow thickness and number values both represent the total number of ticks that moved from a given release plot to the next capture plot during the entire study. The plot labeled “Outside” represents the entire area around our five plots. Most ticks were released and recaptured in plot 3, however we did find ticks that moved from plot 3 to plots 1, 2, 4, and 5, as well as to outside the plot boundaries. Similarly, while most ticks that were released in plot 5 were also recaptured within plot 5, some ticks showed movement from plot 5 to plot 4. The river bordering plot 5 on one slide likely limited movement to outside the plot area.

**Fig. 10.**
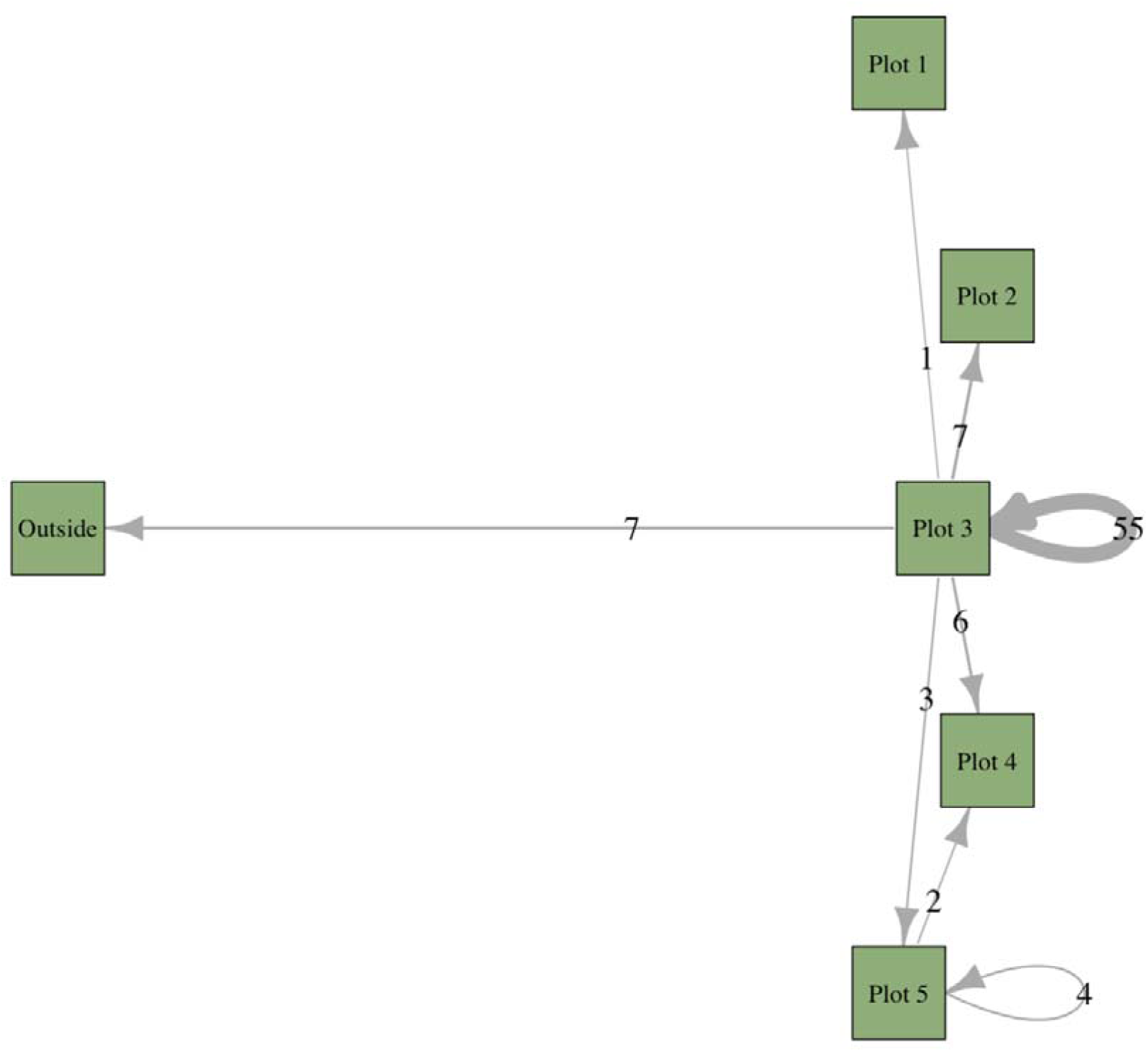
All events of tick movements from release plot to the next capture plot.

## 4 DISCUSSION

In this study, we developed and applied a MRR-based approach to quantify fine-scale dispersal and movement patterns of individual *D. andersoni* ticks. This approach incorporates a simple, low-cost marking strategy that enables unique identification of hundreds of individuals, making large-scale, individual-level movement studies of ixodid ticks feasible under natural field conditions. Using this framework, we observed non-random movement patterns, including directed movement towards grass, vegetation-dependent dispersal distances and rate, and sex-based differences in movement. We also observed relatively high recapture rates, further supporting the feasibility of this approach. Together, these findings highlight the value of this MRR approach for advancing our understanding of tick dispersal in natural systems.

Ticks have been found to exhibit directed movement across heterogeneous landscapes. In a study examining *I. scapularis* released at a woodland–pasture interface, nymphs were disproportionately more likely to disperse into wooded habitats, whereas adults were more frequently recaptured in pastures (Carroll & Schmidtmann, 1996). These contrasting movement patterns likely reflect life stage specific differences in seasonal activity and habitat requirements. Nymphal *I. scapularis* are most active during the summer months, when microclimatic conditions in open pastures may be unfavorable due to higher temperatures and lower humidity. As a result, nymphs may preferentially disperse toward wooded areas, where increased canopy cover provides cooler, more humid conditions. In contrast, adult *I. scapularis*, which are more resistant to desiccation and primarily active in the fall, show a less pronounced preference for pastures. Climatic differences between wooded and pasture habitats are less distinct in the fall, so habitat preferences may instead be guided by other factors, such as host availability or vegetation abundance. Similarly, in this study, we found directed movement towards certain vegetation types. Specifically, we found that host-seeking *D. andersoni* ticks were captured at disproportionately higher rates on grass, whereas ticks were captured at disproportionately lower rates on bare ground, juniper, sapling, and shrub. This preference for grass for host-seeking activity may reflect structural characteristics of the vegetation, including height, blade width, or blade density, ease of movement between the soil and vegetation layers, or higher likelihood of host movement through these areas. Collectively, these patterns underscore the importance of vegetation structure and associated microclimatic conditions in shaping tick movement across heterogeneous landscapes.

Reported tick dispersal distances and rates vary widely across species and studies, including substantial variation also observed within species. Across taxa, reported dispersal distances ranged from 60 cm in *D. reticulatus* to 15.6 m in *A. americanum* (Buczek et al., 2017; Koch & McNew, 1982). Even within *I. scapularis*, estimates of dispersal distance differ markedly across studies; Goddard (1993) found that adult *I. scapularis* do not move laterally to a significant extent (<0.5 m) during questing (Goddard, 1993), while Carroll and Schmidtmann (1996) reported that adult *I. scapularis* dispersed 3.7 m on average and up to 7-8 m overall (Carroll & Schmidtmann, 1996). These reported differences may be due to specific ecological contexts or weather conditions during the study period. Supporting this interpretation, a multi-year study of *D. reticulatus* found that movement rates varied dramatically across study years (Buczek et al., 2017). Interestingly, land management practices (mechanical thinning) took place between study years, indicating that shifting environmental conditions may have a large effect on tick behavior. For example, ticks may disperse longer distances when weather conditions (e.g., temperature, humidity) are more favorable due to lower risk of desiccation and remain more stationary or retreat to the soil layer when conditions are less favorable. Taken together, these patterns indicate that tick dispersal is highly context dependent and responsive to local environmental conditions. Our findings are consistent with this framework; we found that *D. andersoni* ticks dispersed an average of 0.67 meters per day, but that dispersal distance and rate was not constant and instead depended strongly on the vegetation type on which ticks were released, as well as seasonality. Vegetation-specific differences in microclimate and host-seeking suitability may therefore contribute to the substantial variation in dispersal behaviors reported across studies. Additionally, ticks may be more willing to risk desiccation when located at unfavorable host-seeking vegetation in search for more suitable habitats, and more likely to remain stationary when located upon favorable vegetation. Overall, this evidence suggests tick dispersal is a dynamic process shaped by ecological conditions, emphasizing the need to consider how shifts in habitat structure and microclimate may influence tick movement, survival, and host encounter rates.

Among non-nidicolous, ambush-feeding ticks, reported dispersal distances vary by life stage and sex. In a study of *D. reticulatus* ticks, it was found that female ticks dispersed farther than male ticks (Buczek et al., 2017). It has also been shown that adult *I. scapularis* ticks disperse greater distances than nymphs (Carroll & Schmidtmann, 1996). These differences between sex and life stage are likely due to size (with adult ticks being larger than nymphal ticks and adult females being larger than adult males, on average) and preferred host species. Because larval and nymphal ticks typically feed on the same small animal hosts, engorged larvae may be more likely to detach in areas that also support suitable hosts for the nymphal stage. In this study, we were unable to make comparisons between nymphal and adult *D. andersoni* ticks due to the low number of nymphs collected. However, consistent with previous studies, we observed that female *D. andersoni* ticks dispersed farther than male *D. andersoni* ticks. Given these apparent differences in dispersal capacity by life stage and sex, habitat change may differentially affect movement patterns across life stages and sex, with complex consequences for tick population dynamics, including altered mating opportunities.

In MRR studies, recapture rates are shaped by organismal and ecosystem traits and reflect the combined effects of movement, detectability, and loss processes. When recapture rates are low, inference about dispersal patterns becomes more limited because fewer movement events are observed, increasing uncertainty in dispersal estimates. Reduced recapture rates in ticks may result from dispersal beyond the sampling area, retreat into the soil or litter layer, host attachment, or mortality. For these reasons, recapture rates are expected to vary across ecological contexts. Consistent with this expectation, a study comparing *D. variabilis* recapture rates between trails frequently or infrequently visited by hikers, horses, and pets, found that substantially higher recapture probabilities were found on infrequently used trails (Carroll et al., 1991). Weekly recapture probabilities on high use trails ranged from 0.31 to 0.60, whereas low use trails ranged from 0.68 to 0.77, demonstrating the influence of land use on recapture rates. In our study, we recaptured a relatively high proportion (>70%) of ticks at least once during the study period, providing evidence that this MRR approach is feasible for characterizing tick dispersal patterns. Notably, our field site was not located near a trail system and was infrequently visited by recreators, which likely contributed to the high recapture rate we observed. Even so, recapture rates for hard ticks are often relatively high (frequently exceeding 50%) (Bugmyrin & Gorbach, 2022; Carroll & Schmidtmann, 1996) compared to those reported for other disease vectors, particularly mosquitoes, for which recapture rates commonly range from approximately 1-8% depending on the taxa (Guerra et al., 2014). Altogether, the relatively high recapture rates observed in hard ticks enhances the feasibility of MRR approaches for resolving fine-scale dispersal patterns.

A major benefit of this MRR technique is the ability to track individual ticks through space and time, enabling analyses that are not possible with batch marking strategies. Using this approach, we were able to detect vegetation preferences, dispersal distances and rates, and sex-based behavioral differences in *D. andersoni*, demonstrating this method’s utility for characterizing fine-scale tick habitat use and movement. Individual-level marking is particularly valuable for calculating population sizes, quantifying how long ticks quest across a season, how questing duration and movement patterns shift with time of day, weather, or season, and how landscape features influence dispersal. This approach could be applied to make comparisons of dispersal behavior between ambush and hunter tick species or assess how extended questing or increased movement may influence host encounter rates, disease risk, or reproductive success. Finally, when paired with host trapping, this method offers a unique opportunity to quantify host acquisition rates and evaluate how host behavior influences tick movement and recapture. Overall, this method allows us to answer questions that require repeated observations of the same individuals.

Although our approach offers several advantages over other MRR approaches previously used for ticks, there are important limitations to consider. First, using white or light-colored nail polish may make ticks harder to detect on white drag cloths. Nail polish color choices should be selected carefully to maximize contrast with drag cloth color and accommodate color-blind researchers. Second, when a marked tick is not recaptured, the underlying reason remains uncertain. We cannot conclude whether the tick has found a host, died, or is simply not questing at the time of sampling. Repeated observations may reduce this uncertainty by increasing detection probability. Third, marking could potentially impact overall fitness, such as through higher susceptibility to predation. Future controlled studies would help clarify whether and how marking affects tick behavior or fitness. Fourth, this study was limited to a single location and season, highlighting the need for future work across broader spatial and temporal scales. Despite these limitations, MRR remains a powerful tool for studying animal movement and behavior, and will be increasingly important for understanding how climate, disturbances, and land-use changes shape tick populations and TBD risk.

In this study, we used an individual tick marking strategy to understand *D. andersoni* questing vegetation preferences and quantify dispersal behaviors and patterns. We demonstrated the feasibility and effectiveness of this individual marking strategy to address questions that require repeated observations of individual ticks. This method can be applied to other hard tick species to answer questions about individual tick questing patterns, dispersal, and population dynamics across space and time.

## Author Contributions

Elizabeth Hemming-Schroeder conceived the ideas and designed methodology; Sabrina Gobran, Jake Brisnehan, and Jon Wegryn collected the data; Sabrina Gobran and Jake Brisnehan analyzed the results; Sabrina Gobran led the writing of the manuscript; Elizabeth Hemming-Schroeder contributed significantly to the review and editing of the drafts. All authors contributed substantially to the drafts and gave final approval for publication.

## Statement on inclusion

The entirety of this study was undertaken in Colorado. We provided opportunities for intellectual contribution by local researchers by involving undergraduate and graduate researchers from Colorado State University in the design, implementation, and analysis of this study. Trainees contributed to fieldwork, data analyses, and interpretation.

## Acknowledgements

A special thanks to Lawson Dawe for identifying the ticks collected during this study. This work was supported by the Colorado State University One Health Institute, the Infectious Disease Research and Response Training Program (NIH NRSA T32AI162691), and the Rockies and High Plains Vector-borne Diseases Center. The Centers for Disease Control and Prevention, Department of Health and Human Services, provided financial support for this project. The contents are those of the authors. They may not reflect the policies of the Department of Health and Human Services or the US government.

## Data availability statement

Data and code associated with this manuscript is expected to be archived on Zenodo should the manuscript be accepted.

